# Physics-informed neural networks for subject-specific recovery of travelling-wave structure in resting-state fMRI

**DOI:** 10.64898/2026.10.05.756801

**Authors:** Taghipour L. Pouya, Roberto C. Sotero

## Abstract

Resting-state BOLD fluctuations show reproducible spatiotemporal propagation that is not accounted for by conventional zero-lag functional connectivity. Travelling wave analyses based on phase gradients or statistical decomposition have problems when multiple wave generators overlap or when wave origins have to be localized. This work introduces a wave-informed physics-informed neural network that represents the parcellated BOLD signal as a superposition of seven spatially extended damped travelling wave generators, each periodically re-excited at its own characteristic frequency and fixed at initialization to one of the seven canonical Schaefer resting-state networks. A soft anatomical prior derived from each subject’s diffusion-tensor structural connectivity is included to encode the hypothesis that white-matter pathways constrain wave direction. The pipeline is validated on synthetic data by three simulation studies, creating the operating envelope, and then applied to 41 cognitively normal participants of the ADNI 3 cohort. A within-subject ablation showed that mean alignment between recovered wave directions and underlying white-matter geometry increased by a small but statistically detectable amount when using the structural prior (one-sided Wilcoxon signed-rank, *p* = 0.026), with exploratory network-level effects most evident in the visual, dorsal-attention, and salience/ventral-attention networks. The contribution is a network-anchored parametric inverse method returning subject-specific propagation descriptors from short resting-state scans.

## 1 Introduction

Resting-state functional magnetic resonance imaging (rs-fMRI) is widely used to study the brain’s natural functional organization, since spontaneous BOLD fluctuations show structured relationships across distributed regions even in the absence of explicit tasks [1, 2]. Most analyses characterize these relationships using functional connectivity defined through zero-lag correlations. Although this paradigm has been highly valuable for identifying large-scale networks and comparing healthy and diseased populations, it does not capture the temporal ordering of fundamental and natural activity. Increasing evidence suggests that resting-state BOLD signals are not purely synchronous, but instead exhibit reproducible latency structure and propagating organization across space and time [1, 3].

This temporal perspective extends how resting-state dynamics should be interpreted. Mitra et al. showed that intrinsic activity shows stable whole-brain lag structure, with propagation through and across functional networks on characteristic time scales [1]. Hindriks et al. identified travelling BOLD waves in visual cortex linking task-positive and task-negative systems, further supporting the view that meaningful temporal organization exists beyond instant synchrony [3]. At a more local physiological scale, Aquino et al. demonstrated that hemodynamic travelling waves are both reasonable and practically observable in human visual cortex, with an exponential damping envelope referable to vascular dissipation [4]. Studies of quasiperiodic patterns have shown that recurring large-scale spatiotemporal motifs contribute substantially to resting-state organization [5], and subject-specific investigations have shown that propagated intrinsic activity contains specific features, highlighting the importance of studying these dynamics at the single-subject level [8]. Together, these findings point out that propagation-related structure is a fundamental component of the rs-fMRI signal rather than an accidental consequence, and they have motivated broader efforts to model how anatomical organization and dynamical interactions shape whole-brain functional activity [6, 7, 10].

Current methods for analyzing these travelling waves face significant limitations when handling multiple overlapping signals or localizing wave origins. Phase-gradient analysis extracts the instantaneous phase of the BOLD signal to estimate local wave direction and speed [3]; while effective when a locally dominant travelling component is present, overlapping waves can make the resulting phase gradients difficult to interpret. When multiple intrinsic networks generate overlapping waves, the resulting spatial phase gradients can become difficult to interpret, potentially producing apparent singularities or mixed propagation patterns that do not separate the underlying generators. To handle more complex spatiotemporal patterns, other studies employ dimensionality reduction and dynamical systems approaches such as Dynamic Mode Decomposition or complex-valued ICA [11, 12]. These methods successfully extract multiple spatiotemporal components, but they break down the signal into independent statistical modes or linear mathematical approximations rather than distinct physical wave generators, and neither approach explicitly estimates the parametric spatial origins of the underlying waves. Comparatively less attention has been given to mechanistic, inverse-problem frameworks designed to address this superposition problem directly. Physics-informed neural networks (PINNs) [13] have recently been applied to estimate parameters in macroscopic brain-dynamics models [15] and to extract directed functional connectivity from rs-fMRI [16, 17], but the framework has not yet been adapted to isolate and parameterize overlapping travelling waves with explicit physical structure. This work introduces a wave-informed PINN that represents the parcellated BOLD signal as a superposition of seven spatially extended damped travelling-wave generators, each periodically re-excited at its own characteristic frequency and anchored at initialization to one of the seven canonical resting-state networks of the Schaefer parcellation [9]. By embedding the wave equation directly into the loss function, the network is forced to explain the data through the superposition of distinct generators, encouraging strict compatibility between the learned output and physical wave propagation. The framework is additionally regularized by a soft anatomical prior derived from each subject’s diffusion-tensor structural connectivity, encoding the hypothesis that white-matter pathways constrain the spatial direction of hemodynamic propagation. The pipeline is validated on synthetic data through three simulation studies, an initialization-strategy benchmark, a spatiotemporal-resolution sweep, and an SNR-robustness sweep, and then applied to 41 cognitively normal participants of the ADNI 3 cohort. The contribution is dual: a network-anchored pulsed travelling-wave PINN that recovers subject-specific propagation parameters from short rs-fMRI scans, and a within-subject ablation that tests whether structural connectivity carries information about wave geometry over and above what a functional parcellation provides on its own.

## 2 Methods

### 2.1 Data acquisition and preprocessing

The rs-fMRI and diffusion-weighted imaging (DWI) data from 41 cognitively normal participants enrolled in the Alzheimer’s Disease Neuroimaging Initiative phase 3 (ADNI 3) study were analyzed [18]. All 41 included participants were imaged under the same basic acquisition protocol where the rs-fMRI was a 3 T single-band gradient-echo EPI sequence (TR= 3.0 s, 3.4 × 3.4 × 3.4 mm^3^ voxels, 197 raw volumes per session), and DWI was a single-shell acquisition with *b* = 1000 s/mm^2^ comprising 7 *b*=0 and 48 diffusion-weighted volumes. Functional volumes were preprocessed with a custom FSL-based pipeline including volume removal for magnetization steady-state (*N*_*t*_ = 187 retained volumes), slice-timing correction, rigid-body motion correction, MNI152 2 mm normalization through the structural T1, 6 mm FWHM Gaussian smoothing, and nuisance regression of 14 confounds (six motion parameters, their first temporal derivatives, and the mean white-matter and CSF signals). Global signal regression was not applied because it can alter the temporal and spatial structure of resting-state BOLD correlations and remains methodologically debated [20]. Residuals were band-pass filtered between 0.01 and 0.08 Hz with linear detrending. Preprocessed BOLD volumes were parcellated using the Schaefer 100-region atlas with seven-network organization [9], yielding a (*N*_*t*_, *N*_*r*_) = (187, 100) regional time-series matrix per subject; the MNI coordinates of each region’s centroid, **r**_*i*_ ∈ ℝ^3^, were used as the spatial substrate for the wave model. Diffusion volumes were preprocessed with FSL for eddy-current/motion correction and tensor fitting. Whole-brain deterministic tractography was performed in native diffusion space using DIPY [21] within an FA>0.2 white-matter mask. The Schaefer atlas was transformed into each subject’s diffusion space, and streamline counts between atlas regions were used to construct the subject-specific structural connectivity matrix. The matrix was symmetrized, normalized by the geometric mean of regional voxel counts, and scaled by its global maximum to yield 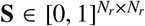. The full preprocessing protocol used here follows the pipeline described in [19].

### 2.2 Travelling-wave forward model

The forward model represents the parcellated BOLD signal as a superposition of *N*_g_ spatially extended damped travelling-wave generators, each periodically re-excited (“pulsed”) at its own characteristic frequency. The structure is motivated by two converging strands of evidence: that hemodynamic responses propagate as spatiotemporal travelling waves with an exponential damping envelope attributable to vascular dissipation [4], and that resting-state BOLD activity exhibits transient, recurrent event-like dynamics rather than purely stationary oscillations [1, 3]. For a generator indexed by g, the contribution to region *i* at time *t* is

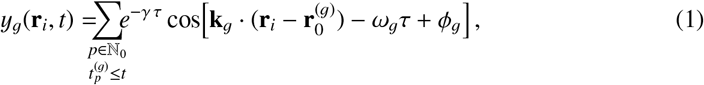

where 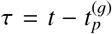 is the time elapsed since the *p*-th pulse-restart event of generator g, with pulse triggers at 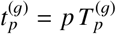 and restart period 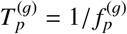. Each generator is therefore characterized by its spatial origin 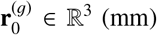, wave vector **k**_g_ ∈ ℝ^3^ (rad/mm) decomposed into a unit direction 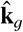 and magnitude |**k**_g_|, angular carrier frequency ω_g_ (rad/s), pulse-restart frequency 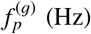, temporal damping rate γ (s^−1^), and carrier phase offset ϕ_g_ (rad). The wave amplitude is hardcoded at unity to remove the well-known amplitude–damping degeneracy of damped oscillator models, and γ is fixed at 0.65 s^−1^, the population mean reported by Aquino et al. [4]. The total forward signal 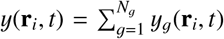 is normalized post hoc to match the empirically preprocessed BOLD: each region is *z*-scored across time and the resulting tensor is divided by its global maximum absolute value, yielding values strictly in [−1, 1].

The number of generators *N*_g_ is fixed at 7, matching the seven canonical resting-state networks of the Schaefer parcellation [9]. Anchoring each generator to a canonical network enables network-stratified analyses on real data, and exploratory work confirmed that data-driven model-order selection criteria (BIC, NMF scree analysis) failed to recover the true generator count on synthetic benchmarks, while the network-anchored (*N*_g_=7) initialization produced consistently stronger parameter recovery.

### 2.3 Physics-informed neural network

The wave model is embedded in a PINN framework [13, 14] that combines flexible data-driven function approximation with explicit physical regularization. The architecture comprises two parallel branches: an LSTM-based encoder *N*_θ_ that reads time and produces a smooth estimate of the regional BOLD signal, and the analytical wave model *W*_Ψ_ from Eq. (1) parameterized by 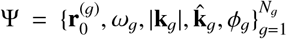. The time variable is mapped to a 6-dimensional feature vector combining a normalized time index with five sinusoidal basis functions linearly spaced over the BOLD band [0.01, 0.10] Hz, accelerating convergence on the oscillatory BOLD signal. The LSTM has two stacked recurrent layers with 128 hidden units and dropout 0.1, followed by a linear projection to *N*_*r*_=100 regional channels.

The total training objective is a weighted sum of three terms,

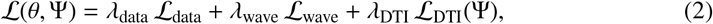

where 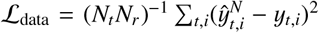 is the standard mean-squared-error data fidelity term that drives *N*_θ_ to reconstruct the preprocessed BOLD; 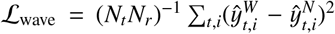 is the physics consistency term that constrains the wave model’s output to match the LSTM’s smooth signal estimate; and ℒ_DTI_ is the diffusion-tensor structural prior detailed below. Weights are fixed at λ_data_ = λ_wave_ = 1.0 and λ_DTI_ = 0.05, an order of magnitude smaller so that the structural prior acts as a soft regularizer. The asymmetric coupling, training the wave model against **ŷ**^*N*^ rather than directly against the raw BOLD, estimates the wave parameters from a denoised signal estimate, mitigating noise-induced bias in parameter recovery; the conditioning is bidirectional through joint optimization, since the wave loss likewise regularizes _θ_ toward travelling-wave–consistent solutions.

All physical wave parameters are bounded by sigmoid reparameterization with bounds ω ∈ [0.06, 0.63] rad/s, |**k**| ∈ [0.03, 0.15] rad/mm, ϕ ∈ [−π, π], and **r**_0_ within the atlas extent. The wave-vector direction 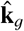 is parameterized as a free ℝ^3^ vector divided by its Euclidean norm, and the phase ϕ_g_ uses a tanh-bounded reparameterization. Training proceeds in two stages: a 100-epoch warm-up in which only the LSTM parameters are optimized against *ℒ*_data_, and a 500-epoch joint phase under the composite loss of Eq. (2). Both stages use Adam, with learning rate 10^−3^ for the warm-up and 5 × 10^−4^ for the joint stage, and gradient norm clipped at unity. The total training budget is ~ 8 s on an Apple M5 GPU per subject.

### 2.4 Initialization

To break the variation symmetry of unrestricted multi-generator fitting and to inject prior anatomical knowledge, the seven generators are initialized in a network-anchored, data-driven manner. For each Schaefer canonical resting-state network, one anatomical region *i*_g_ is selected at random from within the regions belonging to that network, and its centroid is used as the initial spatial origin 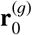. From the Hilbert analytic signal of the preprocessed regional BOLD at the anchor region, the pulse-restart frequency 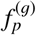 is initialized as the spectral peak of the envelope time series within the BOLD band, and the carrier angular frequency ω_g_ is initialized as the peak of the raw BOLD power spectrum within the same band, with a ±0.005 Hz exclusion around 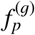 to ensure pulse and carrier frequencies remain spectrally distinct. The wave vector **k**_g_ is initialized via phase-coordinate regression: relative phase offsets across regions, computed as the circular mean of the relative instantaneous phase, are fit by least squares against the spatial displacements from the anchor centroid, and the resulting estimate is decomposed into a unit direction and a magnitude. The pulse frequency 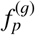 is held *fixed* at its data-derived value throughout optimization, since the strong dependence of the pulse-restart structure on its empirical estimate makes refitting destabilizing on noisy data; the phase ϕ_g_ is initialized uniformly at random in [−π, π]. The trainable parameter set 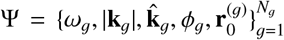 contains 8 effective degrees of freedom per generator, totaling 56 free parameters across the seven generators.

### 2.5 Diffusion-tensor structural prior

The hypothesis that white-matter pathways constrain the spatial direction of hemodynamic travelling waves is operationalized as a soft regularization term on the wave-vector orientations. For generator g with origin 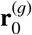, let *n*_g_ denote the atlas region whose centroid is nearest to that origin, and let 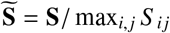 be the SC matrix rescaled so that its maximum entry is unity. The unit vector from the origin to each atlas region *j* is 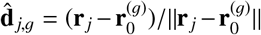. The DTI penalty for one generator is the SC-weighted mean of the angular misalignment between the recovered direction 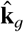 and the radial unit vectors 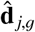 summed across the cortex,

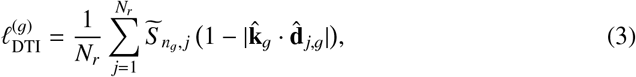

and the overall structural-prior loss averages over the seven generators, 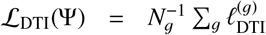. The absolute cosine | · | enforces axial rather than vector alignment, the appropriate symmetry for white-matter tracts. The ablation comparison between λ_DTI_ = 0.05 and λ_DTI_ = 0 provides the experimental test of the white-matter-constraint hypothesis on real ADNI data.

### 2.6 Validation

The framework was validated on synthetic BOLD generated by the forward model. Ground-truth wave parameters were drawn uniformly from the optimization bounds, with γ sampled from the Aquino range [0.5, 0.9] s^−1^ and origins sampled within [−50, 50]^3^ mm of the parcellation centroid; synthetic structural connectivity matrices used in DTI runs were generated as block-diagonal patterns aligned with each wave’s 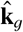 as a positive control. Independent Gaussian noise with standard deviation σ was added at the regional level before normalization. Recovery was reported on four parameters: *f*_*p*_, ω, propagation direction 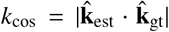, and origin localization error. Three studies were run: an initialization-strategy benchmark, a spatiotemporal-resolution sweep, and an SNR-robustness sweep. The initialization-strategy benchmark compares the network-anchored pipeline against a free-*N*, NMF-initialized baseline across *N*_true_ ∈ {1, 2, 3, 5}, σ ∈ {0.0, 0.1, 0.2}, and 10 seeds per condition, with and without the DTI constraint. The spatiotemporal-resolution sweep uses seven ground-truth waves, five fixed background generators and two target generators whose origin separation Δ*r* (10–100 mm) and pulse-frequency separation Δ *f*_*p*_ (0–50 mHz) are systematically varied along a 7 × 7 grid. The SNR-robustness sweep uses *N*_true_ = 3 across 16 noise levels spanning σ ∈ [0, 3.0] in unit-normalized signal space, with 10 seeds per level.

### 2.7 Real-data application and statistical analysis

The pipeline was applied to all *N*_sub_ = 41 cognitively normal participants. Each subject was processed twice with identical hyperparameters, random seeds, and initialization: once with the structural prior enabled (λ_DTI_ = 0.05) and once with it disabled (λ_DTI_ = 0). The two paired runs share the same network-anchored origins, the same data-driven {*f*_*p*_, ω, **k**} initializations, and the same LSTM warm-up, so any difference in recovered parameters is attributable to the DTI term and not to optimization noise.

For each participant, a local SC-weighted direction was computed at every parcel as the unit vector pointing along the strongest white-matter connectivity from that parcel, given by the SC-weighted sum of unit vectors to all other parcels and renormalized. For each recovered generator g, the alignment with local white-matter geometry is 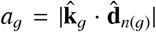, where *n*(g) is the parcel nearest to the recovered origin. The absolute value reflects the axial balance of white-matter tracts and the sign invariance of Eq. (1) under simultaneous flips of 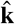 and ϕ.

The directional hypothesis, that the DTI-constrained run yields higher alignment with white-matter structure than the unconstrained run, was tested by summarizing each participant by the mean alignment across that participant’s seven generators 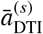 and 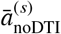 and applying a one-sided paired Wilcoxon signed-rank test of the alternative 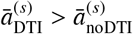 across the *N*_sub_ = 41 participants. The Wilcoxon test was preferred over a paired *t*-test because the alignment score is bounded and its small-sample distribution is not assumed Gaussian; the one-sided alternative is justified by the directional nature of the mentioned hypothesis. To localize where the effect originates, the same one-sided paired Wilcoxon test was applied within each Schaefer 7-network individually, using only that network’s generator from each participant. Per-network *p*-values are reported uncorrected; with the seven networks defined a priori by the model architecture rather than chosen post hoc, the partition is treated as a planned exploratory decomposition rather than as an independent multiplicity-corrected test.

## 3 Results

The initialization-strategy benchmark compared the network-anchored pipeline against a free-*N*, NMF-initialized baseline. Aggregate recovery metrics across the twelve scenario conditions and ten seeds per condition are summarized in Table 1: the network-anchored pipeline outperformed the baseline on every metric, lifting mean direction cosine from 0.667 to 0.750, pulse-frequency recovery from 74% to 83% of wave instances within one standard deviation of ground truth, and reducing mean origin localization error by roughly six millimeters. A second pattern emerged in the structural-prior contrast: with the free-*N* baseline, enabling the DTI loss raised *k*_cos_ in all twelve conditions, with a mean lift of Δ*k*_cos_ = +0.068. With the network-anchored pipeline, the same contrast averaged only +0.018 and was zero or slightly negative in five of twelve conditions. When seven generators are spread across the seven Schaefer networks at initialization, the optimization already starts close to a good local minimum, and the synthetic block-diagonal structural prior has little additional spatial information to inject. This is a ceiling effect rather than evidence against the prior; the real-data analysis tests whether the same ceiling holds when the structural connectivity reflects genuine biological tract anatomy.

**Table 1:** Aggregate parameter recovery in the initialization-strategy benchmark.

| Metric | Free-$N$ baseline | Network-anchored |
| --- | --- | --- |
| $f_p$ recovery rate (within $\pm 1$ SD) | 74% | **83%** |
| Mean $k_{\cos}$ | 0.667 | **0.750** |
| Mean origin error (mm) | 80.2 | **74.4** |
| Mean $\Delta k_{\cos}$ (DTI on – off) | +0.068 | +0.018 |

The network-anchored initialization is a stronger baseline for fMRI travelling-wave recovery than a free-*N* scheme that selects origins from peaks of NMF spatial maps. The improvement is meaningful but not dramatic. These numbers describe a method that works well enough to be useful as a population-level descriptor of wave geometry but not well enough to localize individual generators with surgical precision. These results support interpreting the method as a regional, network-scale descriptor of propagation geometry rather than as a precise localization tool for individual wave sources.

The structural prior contributes little additional information *on synthetic data* once the network-anchored initialization is in place: the mean DTI lift on synthetic data was +0.018 in *k*_cos_, compared to +0.068 for the free-*N* baseline. This is a ceiling effect rather than a failure: the network anchoring already places generators where the brain’s modular organization predicts they should sit, and the synthetic block-diagonal structural connectivity matrices used in simulation provide little orthogonal information beyond that anchoring. The natural next question is whether a real diffusion-derived structural connectome, with its measured anisotropies and biological tract geometry, behaves similarly; the cohort analysis suggests that it does not.

The spatiotemporal-resolution sweep probed the spatiotemporal resolution of the pipeline by varying the spatial separation Δ*r* and pulse-frequency separation Δ *f*_*p*_ between two target generators in the presence of five fixed background generators (Figure 1). Three patterns emerge from the heatmaps. Pulse-frequency error grows with Δ *f*_*p*_ rather than with Δ*r*: along the Δ *f*_*p*_ = 0 row, the empirically fixed *f*_*p*_ at the initialization seed is correct for both targets and the model recovers *f*_*p*_ with errors below 20%, but as the targets’ true frequencies separate, a single empirical seed can no longer represent both, and errors climb to 40-80% at the largest Δ *f*_*p*_. The carrier oscillation ω is essentially unidentifiable across the entire grid, with errors ranging from 75% to over 160% and no clean dependence on either axis; this is consistent with the known difficulty of separating multiple narrowband oscillators within the BOLD passband and is reported transparently rather than masked. Direction recovery *k*_cos_ and origin error *r*_0_ (not shown) were roughly flat across the grid in this five-background regime, with *k*_cos_ in 0.52–0.73 and *r*_0_ in 67–99 mm, neither showed a clean monotonic improvement with Δ*r* at the parcel scale used here, indicating that with five background generators present, the marginal contribution of any individual target is bounded by how well the optimizer can untangle the overall scene rather than by how far apart the two targets sit.

**Figure 1:**
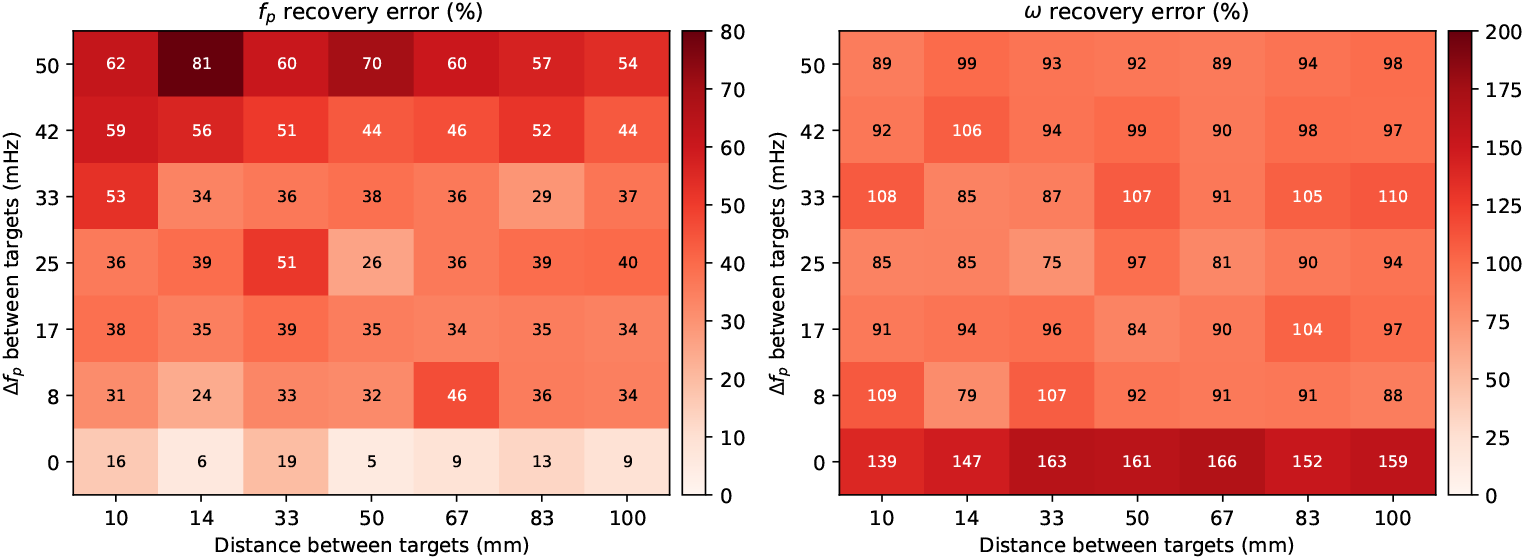
Spatiotemporal-resolution sweep. Heatmaps of (left) *f*_*p*_ recovery error and (right) ω recovery error for the two target generators as a function of spatial separation Δ*r* (x-axis) and pulse-frequency separation Δ *f*_*p*_ (y-axis), with five fixed background generators present. *f*_*p*_ error is dominated by the Δ *f*_*p*_ axis rather than by Δ*r*; ω is broadly unidentifiable.

The SNR-robustness sweep characterized recovery as a function of signal-to-noise ratio (Figure 2). Recovery was remarkably stable across the realistic fMRI SNR range. The direction cosine *k*_cos_ stayed within [0.71, 0.81] from noiseless conditions down to roughly −6 dB SNR. A counter-intuitive but real observation is that the ω error decreased with increasing noise, from roughly 65% in the noiseless case to around 35% at σ = 0.45. This pattern is consistent with stochastic regularization: in a strongly constrained optimization landscape (fixed γ, *A* = 1, fixed *f*_*p*_, network-anchored origins), small amounts of noise help the optimizer escape shallow local minima rather than degrading the fit. Origin localization hovered near 70–80 mm and *f*_*p*_ error stayed near 40–50 % across the tested span (not shown). Rather than presenting a sharp degradation curve as initially expected, this study demonstrates that the pipeline is noise-robust within the operating regime relevant for resting-state BOLD.

**Figure 2:**
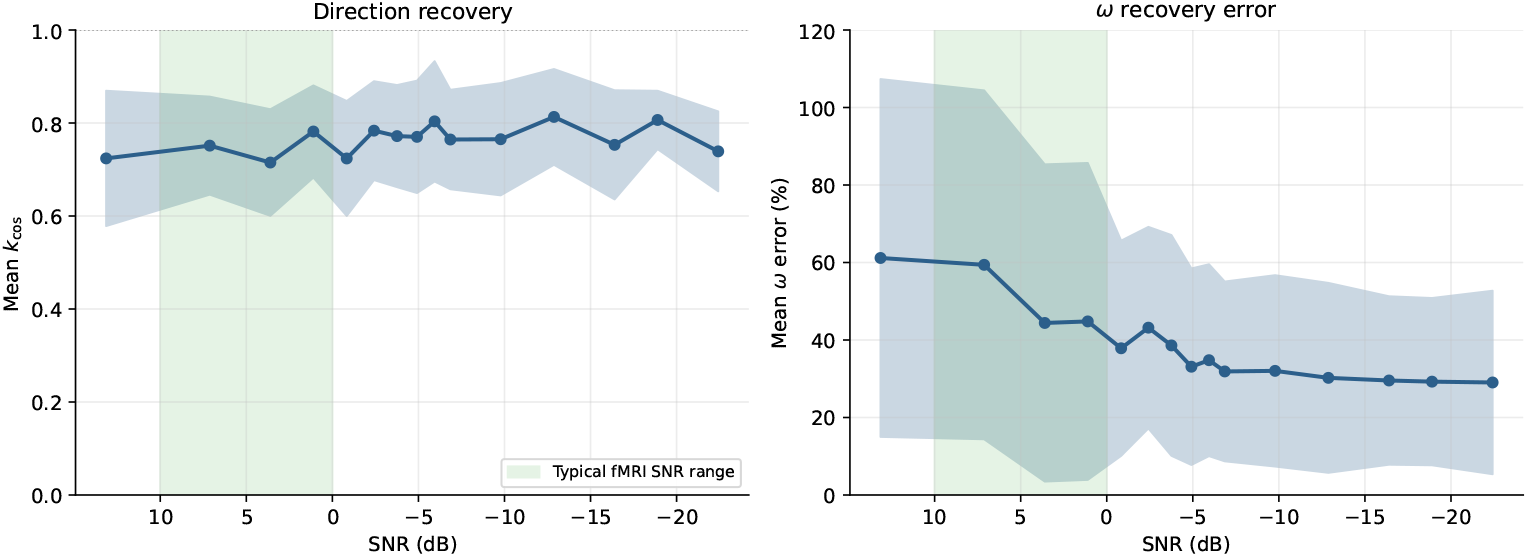
SNR-robustness sweep. Recovery metrics as a function of SNR (decibels, x-axis reversed so easier conditions are on the left). Left: *k*_cos_ remains in [0.71, 0.81] across the entire tested SNR span. Right: ω error decreases with added noise, consistent with stochastic regularization. Shaded bands show ±1 SD across ten seeds per noise level.

Turning to the cohort analysis, the pipeline was applied to all forty-one cognitively normal ADNI 3 subjects, with each subject processed twice under identical seeds and hyperparameters, differing only in whether the structural prior was active. The contrast of interest was the within-subject change in mean structural alignment 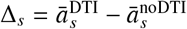, with hypothesis predicting Δ_*s*_ > 0. A one-sided Wilcoxon signed-rank test on the forty-one paired observations rejected the null hypothesis of zero median difference, with *p* = 0.0264 and a median paired increase of Δ_*s*_ = +0.028 in the structural-alignment score (Figure 3, Panel A). The effect is small in absolute terms but consistent in sign across the cohort, and it cannot be linked to differences in initialization, atlas, or seed because every other element of the pipeline was held identical between the paired runs.

**Figure 3:**
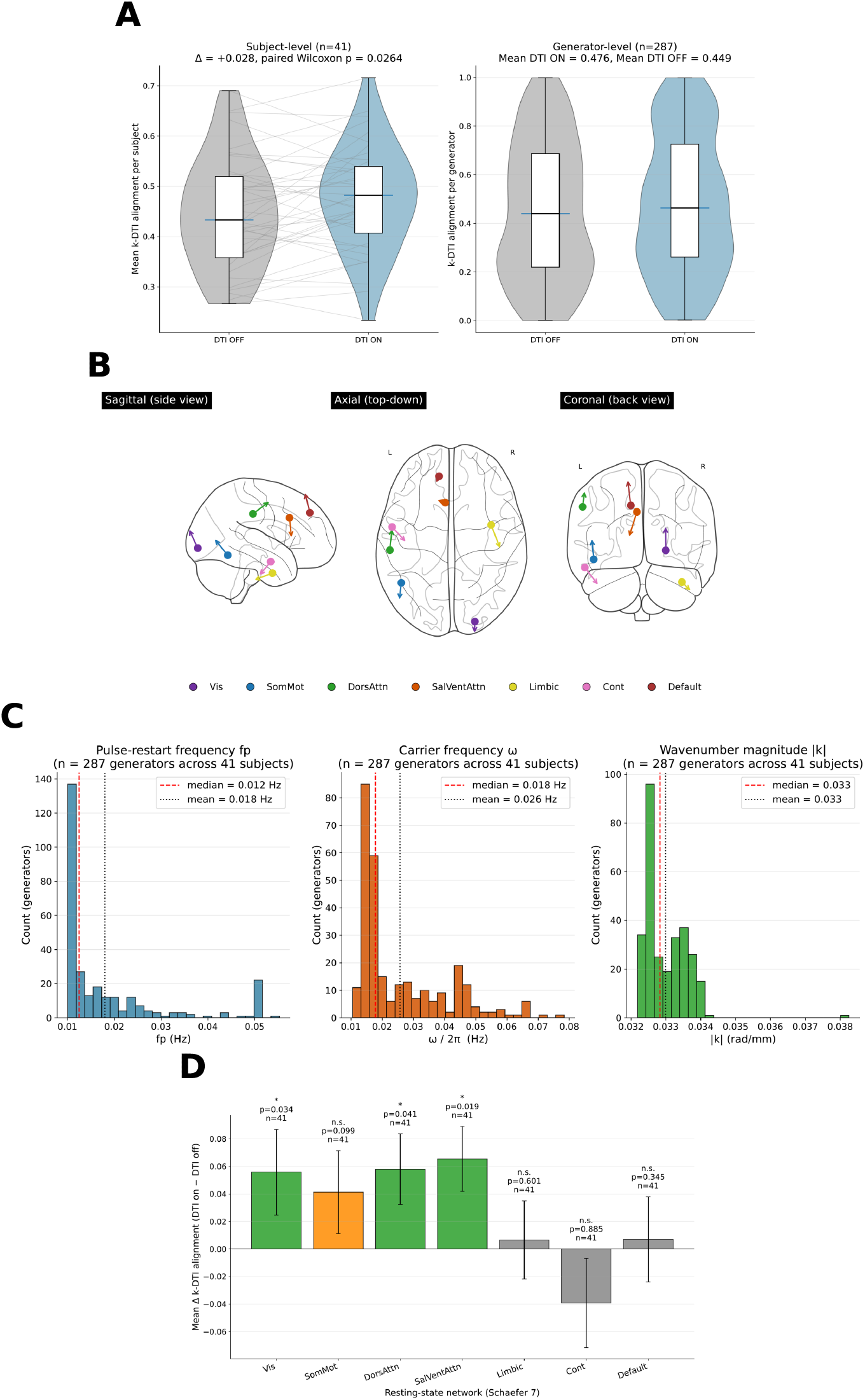
Real-data analysis on the ADNI 3 cohort (*N* = 41). (A) Subject-level structural alignment with and without the DTI prior. (B) Recovered cortical generator origins and directions. (C) Distributions of recovered *f*_*p*_ and ω across 287 subject-generator pairs. (D) Network-level structural-alignment effects.

The recovered origins and propagation vectors distribute across the cortex rather than collapsing onto a single hub (Figure 3, Panel B). The propagation vectors span the full range of orientations rather than aligning trivially with any single anatomical axis, this pattern is consistent with biologically plausible recovery and argues against the pathological failure mode in which the optimizer shrinks all generators to a single point or aligns all directions with a fixed coordinate axis. The cohort-level distributions of recovered *f*_*p*_ and ω values, pooled across the 41 × 7 = 287 subject-generator pairs (Panel C), are smooth and unimodal within the physiologically plausible range, with no evidence of mass piling up on the bound boundaries, the kind of population-level convergence one would hope to see from a method intended for cohort-scale clinical use.

Partitioning the contrast by Schaefer 7-network reveals that the cohort-level effect is driven primarily by a subset of the networks (Figure 3, Panel D). Three of the seven networks reach the conventional significance threshold individually under the one-sided Wilcoxon test: visual (Vis, *p* = 0.034), dorsal attention (DorsAttn, *p* = 0.041), and salience/ventral attention (SalVentAttn, *p* = 0.019); somatomotor (SomMot) shows a trend (*p* = 0.099), and the remaining three networks (Limbic, Cont, Default) are non-significant. One possible interpretation is that the structural prior is more informative in sensory and attentional systems, where dominant white-matter pathways may impose stronger directional constraints on propagation geometry; in higher-order associative networks, where tract anatomy may be more distributed or where generators are already well-localized by the network-anchored initialization, the marginal information added by the structural prior appears smaller. This decomposition is exploratory and is reported descriptively rather than as an independently corrected test. The difference between the synthetic and real-data prior effects is an important observation, because it suggests that the empirical diffusion-derived connectome contains geometric information not captured by the simplified synthetic block-diagonal connectomes. The contrast was constructed so that nothing differed between paired runs except whether the structural-alignment loss term contributed to the objective, eliminating between-subject confounds by construction. The fact that the effect appears on real brains while being largely absent on synthetic data indicates that real diffusion-derived structural connectivity carries information about the geometry of resting-state wave propagation that synthetic block-diagonal connectomes cannot replicate. For example, visual and attention-related systems are supported by relatively organized projection and association pathways, whereas higher-order associative networks may have more distributed connectivity patterns or may already be sufficiently constrained by the network-anchored initialization. This pattern is consistent with prior work showing that lag structure in resting-state BOLD reflects axonal pathway organization [1], and with broader literature linking structural connectivity to the spatiotemporal organization of intrinsic brain dynamics [4].

Taken together, these results support a single, modest claim: a network-anchored physics-informed pipeline can recover travelling-wave propagation directions from resting-state BOLD with reasonable robustness across SNR and scenario configurations, and a soft diffusion-tensor structural prior, while contributing little additional information on synthetic positive controls, produces a small but reproducible improvement in within-subject structural alignment on real cognitively normal brains, concentrated in the sensory-attentional systems whose white-matter anatomy is most strongly directional.

## 4 Conclusion

This study introduced a physics-informed neural network for recovering pulsed travelling-wave structure from rs-fMRI, anchored to the seven Schaefer networks at initialization and softly regularized by diffusion-tensor structural connectivity. Three findings stand out, and each carries a corresponding warning that should shape how the work is read and how it is extended.

The cohort is small and homogeneous: forty-one cognitively normal subjects from a single ADNI 3 acquisition protocol. Replication in larger and more diverse cohorts is essential before any claim about clinical relevance can be made, and the seven-network anchoring, the fixed value of γ, and the assumption of stationarity over the imaging duration are all defensible but testable modeling commitments that future work could relax.

The natural next step is to test whether the structural-alignment effect generalizes across clinical populations and whether the recovered wave parameters track disease-relevant variation in connectivity; the ADNI 3 cohort already includes mild cognitive impairment and Alzheimer’s disease subjects, and applying the same pipeline to those cohorts is a direct follow-on study. In its present form, the contribution is methodological rather than clinical: a working pipeline, a calibrated operating envelope, and a within-subject demonstration that diffusion-derived structural connectivity carries information about the geometry of fMRI travelling waves over and above what a network parcellation provides on its own.

## References

[1] Mitra, A. Z. Snyder, C. D. Hacker, and M. E. Raichle, “Lag structure in resting-state fMRI,” Journal of Neurophysiology 111(11), 2374–2391 (2014). 10.1152/jn.00804.2013

[2] Lennartz, J. Schiefer, S. Rotter, J. Hennig, and P. LeVan, “Sparse estimation of resting-state effective connectivity from fMRI cross-spectra,” Frontiers in Neuroscience 12, 287 (2018). 10.3389/fnins.2018.00287

[3] R. Hindriks, R. Mantini, N. Gravel, and G. Deco, “Latency analysis of resting-state BOLD-fMRI reveals traveling waves in visual cortex linking task-positive and task-negative networks,” NeuroImage 200, 259–274 (2019). 10.1016/j.neuroimage.2019.06.007

[4] K. M. Aquino, M. M. Schira, P. A. Robinson, P. M. Drysdale, and M. Breakspear, “Hemodynamic traveling waves in human visual cortex,” PLoS Computational Biology 8(3), e1002435 (2012). 10.1371/journal.pcbi.1002435

[5] G. J. Thompson, W.-J. Pan, M. E. Magnuson, D. Jaeger, and S. D. Keilholz, “Quasi-periodic patterns (QPP): Large-scale dynamics in resting state fMRI that correlate with local infraslow electrical activity,” NeuroImage 84, 1018–1031 (2014). 10.1016/j.neuroimage.2013.09.029

[6] M. Gilson, N. E. Kouvaris, G. Deco, J.-F. Mangin, C. Poupon, S. Lefranc, D. Riviere, and G. Zamora-Lopez, “Network analysis of whole-brain fMRI dynamics: A new framework based on dynamic communicability,” NeuroImage 201, 116007 (2019). 10.1016/j.neuroimage.2019.116007

[7] M. Pope, C. Seguin, T. F. Varley, J. Faskowitz, and O. Sporns, “Co-evolving dynamics and topology in a coupled oscillator model of resting brain function,” NeuroImage 277, 120266 (2023). 10.1016/j.neuroimage.2023.120266

[8] R. V. Raut, A. Mitra, S. Marek, M. Ortega, A. Z. Snyder, A. Tanenbaum, T. O. Laumann, N. U. F. Dosenbach, and M. E. Raichle, “Organization of propagated intrinsic brain activity in individual humans,” Cerebral Cortex 30(3), 1716–1734 (2020). 10.1093/cercor/bhz198

[9] A. Schaefer, R. Kong, E. M. Gordon, T. O. Laumann, X.-N. Zuo, A. J. Holmes, S. B. Eickhoff, and B. T. T. Yeo, “Local-global parcellation of the human cerebral cortex from intrinsic functional connectivity MRI,” Cerebral Cortex 28(9), 3095–3114 (2018). 10.1093/cercor/bhx179

[10] J. A. Roberts, L. L. Gollo, R. G. Abeysuriya, G. Roberts, P. B. Mitchell, M. W. Woolrich, and M. Breakspear, “Metastable brain waves,” Nature Communications 10, 1056 (2019). 10.1038/s41467-019-08999-0

[11] T. Bolt, J. S. Nomi, D. Bzdok, J. A. Salas, T. Ito, L. Q. Uddin, and S. D. Keilholz, “A parsimonious description of global functional brain organization in three spatiotemporal patterns,” Nature Neuroscience 25, 1093–1103 (2022). 10.1038/s41593-022-01118-1

[12] J. Casorso, X. Kong, W. Chi, D. Van De Ville, B. T. T. Yeo, and R. Liégeois, “Dynamic mode decomposition of resting-state and task fMRI,” NeuroImage 194, 42–54 (2019). 10.1016/j.neuroimage.2019.03.019

[13] M. Raissi, P. Perdikaris, and G. E. Karniadakis, “Physics-informed neural networks: A deep learning framework for solving forward and inverse problems involving nonlinear partial differential equations,” Journal of Computational Physics 378, 686–707 (2019). 10.1016/j.jcp.2018.10.045

[14] G. E. Karniadakis, I. G. Kevrekidis, L. Lu, P. Perdikaris, S. Wang, and L. Yang, “Physics-informed machine learning,” Nature ℝeviews Physics 3(6), 422–440 (2021). 10.1038/s42254-021-00314-5

[15] R. C. Sotero and J. M. Sanchez-Bornot, “Estimating the Excitatory-Inhibitory Balance from Electrocorticography Data using Physics-Informed Neural Networks,” Proceedings of the 11th International Conference on Bioinformatics ℝesearch and Applications, 113–118 (2025). 10.1145/3700666.3700701

[16] R. C. Sotero and J. M. Sanchez-Bornot, “Directed fMRI-based Functional Connectivity Estimation using Physics-Informed Neural Networks,” Proceedings of the 2024 2nd International Conference on Machine Learning and Pattern ℝecognition, 53–58 (2024).

[17] R. C. Sotero, J. M. Sanchez-Bornot, and I. Shaharabi-Farahani, “Parameter Estimation in Brain Dynamics Models from Resting-State fMRI Data using Physics-Informed Neural Networks,” 2024 8th International Conference on Biomedical Engineering and Applications (ICBEA), 73–79 (2024). 10.1109/ICBEA62825.2024.00022

[18] M. W. Weiner, D. P. Veitch, P. S. Aisen, L. A. Beckett, N. J. Cairns, R. C. Green, D. Harvey, C. R. Jack Jr., W. Jagust, J. C. Morris, R. C. Petersen, A. J. Saykin, L. Shaw, A. W. Toga, and J. Q. Trojanowski, “The Alzheimer’s Disease Neuroimaging Initiative 3: Continued innovation for clinical trial improvement,” Alzheimer’s & Dementia 13(5), 561–571 (2017). 10.1016/j.jalz.2016.10.006

[19] Y. Iturria-Medina, F. M. Carbonell, R. C. Sotero, F. Chouinard-Decorte, and A. C. Evans, “Multifactorial causal model of brain (dis)organization and therapeutic intervention: Application to Alzheimer’s disease,” NeuroImage 152, 60–77 (2017). 10.1016/j.neuroimage.2017.02.058

[20] K. Murphy and M. D. Fox, “Towards a consensus regarding global signal regression for resting state functional MRI,” NeuroImage 154, 169–173 (2017). 10.1016/j.neuroimage.2016.11.052

[21] E. Garyfallidis, M. Brett, B. Amirbekian, A. Rokem, S. van der Walt, M. Descoteaux, I. Nimmo-Smith, and Dipy Contributors, “Dipy, a library for the analysis of diffusion MRI data,” Frontiers in Neuroinformatics 8, 8 (2014). 10.3389/fninf.2014.00008

